# Fast, accurate local ancestry inference with FLARE

**DOI:** 10.1101/2022.08.02.502540

**Authors:** Sharon R. Browning, Ryan K. Waples, Brian L. Browning

**Affiliations:** Department of Biostatistics, University of Washington, Seattle, WA; Division of Medical Genetics, Department of Medicine, University of Washington, Seattle, WA

**Author notes:** Corresponding authors (SRB), (BLB).

## Abstract

Local ancestry is the source ancestry at each point in the genome of an admixed individual. Inferred local ancestry is used for admixture mapping and population genetic analyses. We present FLARE (<u>F</u>ast <u>L</u>ocal <u>A</u>ncest<u>r</u>y <u>E</u>stimation), a new method for local ancestry inference. FLARE achieves high accuracy through the use of an extended Li and Stephens model, and it achieves exceptional computational performance through incorporation of computational techniques developed for genotype imputation. Memory requirements are reduced through on-the-fly compression of reference haplotypes and stored checkpoints. Computation time is reduced through the use of composite reference haplotypes. These techniques allow FLARE to scale to data sets with hundreds of thousands of sequenced individuals and to provide superior accuracy on large-scale data. FLARE is freely available at https://github.com/browning-lab/flare.

## Introduction

All humans are admixtures of various historical source populations. This admixture has occurred across a range of timescales, from the recent intercontinental admixture in African Americans and Hispanics, to the ancient admixture with Neanderthals that occurred when modern humans migrated out of Africa around 50,000 years ago.

Local ancestry is the source ancestry of an individual’s chromosomes at each point in the genome. Local ancestry can be inferred on cross-continental admixtures for recently admixed groups, such as admixed populations in the Americas which have admixed ancestry deriving from indigenous Americans, West Africans, and Western Europeans. With sensitive methods, local ancestry of recent within-continental admixtures can also be inferred.^1^

Inferred local ancestry is required for admixture mapping. Admixture mapping tests for association between local ancestry and phenotype, and provides a complementary approach to genome-wide association testing in admixed populations.^2; 3^ Local ancestry can act as a proxy for a variant that is not well captured by the available SNParray or sequencing data, such as a structural variant that is difficult to genotype accurately.

Once a variant is found to be associated with a phenotype, local ancestry can be used to investigate the ancestral origin of an allele. For example, in US-based Hispanics an Amerindian-specific variant of the *ACTN1* gene is associated with platelet count in US-based Hispanics,^4^ and an Amerindian-specific variant of the *BCL2L11* gene is associated with urine albumin-to-creatinine ratio.^5^ In African Americans an African-specific variant is associated with kidney disease.^6^ Identification of the ancestry of disease-associated variants is helpful for understanding and addressing disparities in disease rates.^7^

Local ancestry is also useful for population genetics analyses. Local ancestry segments are used to infer demographic history, including the timing of admixture,^8^ the identity of source populations,^8; 9^ and the effective size of ancestral populations.^10; 11^ Local ancestry can be used for recombination rate inference^12; 13^ because changes in ancestry along an individual’s genome represent crossovers that have occurred since admixture. Genomic regions with local ancestry proportions that deviate from the genome-wide average can signal post-admixture selection.^14; 15^

Increasing amounts of high-coverage whole genome sequence data are available from diverse and admixed populations.^16–18^ This presents opportunities, because substantially increasing the number of reference individuals increases the accuracy of local ancestry inference, particularly in resolving within-continent ancestries. However, larger reference panels also increase the computational burden.

Given these opportunities and challenges, we developed FLARE, which is based on the Li and Stephens model for haplotype frequencies^19^ and follows in the footsteps of the HAPMIX and MOSAIC local ancestry inference methods.^1; 20^ The Li and Stephens model has been widely used for genotype phasing and imputation^21–25^ because it provides high accuracy and it can be combined with powerful computational optimizations.^21–27^ Its computation time is linear in the number of genetic markers, and after optimization its computation time is approximately linear in the number of individuals.^22; 25^ Extending this model to incorporate ancestry extends these advantages to local ancestry inference.

HAPMIX pioneered the application of the Li-and-Stephens model to local ancestry inference. We do not compare FLARE with HAPMIX because HAPMIX is limited to two ancestries. Instead, we compare FLARE with MOSAIC, which is a recent method based on the Li and Stephens model that allows for an arbitrary number of ancestries and unknown relationships between reference panels and ancestry. We show that FLARE has better computational performance and accuracy than MOSAIC in our simulation scenarios.

Other frameworks for inferring local ancestry are possible. One of the most popular methods for local ancestry inference that does not use the Li and Stephens model is RFMix. Rather than utilizing a generative model of haplotype frequencies, RFMix is discriminative and employs a conditional random field.^28^ We compare our method to RFMix and show that FLARE is superior in accuracy and computational performance.

FLARE incorporates several computational techniques which allow it to scale to enormous data sets while maintaining high accuracy. FLARE performs on-the-fly compression of reference haplotypes and stores checkpoints when calculating probabilities to reduce memory requirements. FLARE constructs composite reference haplotypes to reduce computation time. These techniques are described in Methods.

FLARE can analyze both SNP-array and whole genome sequence data. Most existing local ancestry inference methods were designed for SNP-array data. For example, the MOSAIC method was tested using only SNP-array data, and we found that modifications to the program parameters were necessary when analyzing sequence data. FLARE can perform local ancestry inference on sequence data without the information loss that would result from substantial marker thinning.

## Methods

### Hidden Markov Model

FLARE uses a hidden Markov model (HMM). The input data are phased reference haplotypes and phased admixed haplotypes. We use the reference haplotypes to infer local ancestry in one admixed haplotype at a time. Each target admixed haplotype is modelled as an imperfect mosaic of reference haplotypes.^19; 20^ For a target haplotype, the unobserved state *S_m_* = (*i, h*) at marker *m* is comprised of the target haplotype’s ancestry, *i*, at that position and the donor reference haplotype, *h*, whose alleles are being copied at that position.

We assume that there are *A* ancestries contributing to the admixed genomes, and that these ancestries are represented by the reference haplotypes. The reference data consist of *J* panels. In our analyses each reference panel is associated with a single ancestry and *A* = *J*. Our algorithm for estimating model parameters (see Supplementary Methods 3) assumes this one-to-one matching, but the remainder of the methodology does not require a one-to-one matching of reference panels and ancestries, and one could have *J* < *A, J* = *A*, or *J* > *A*.

The number of haplotypes in the *j*-th reference panel is denoted *n_j_*. The total number of reference haplotypes is *N* = ∑_*j*_*n_j_*. We write *p_ij_* for the probability that the donor haplotype is from reference panel *j* when the target haplotype is from ancestry *i*.

State transitions between two adjacent marker positions can occur due to crossover events. Crossover events that occur after admixture can change both the ancestry state, *i*, and the reference haplotype *h*. Crossover events that occur prior to admixture do not change the ancestral state but can change the reference haplotype *h*. We model this second class of crossover events using an ancestry-specific switch rate *ρ_i_*. Ancestry-specific switch rates allow each ancestry to have a different effective population size and a different number of reference haplotypes.

The parameter ***μ*** is a vector of length *A* giving the overall ancestry proportions of the admixed samples. The component *μ_i_* is the prior probability that an arbitrary position in the genome is derived from ancestry *i*. Ancestry probabilities sum to one, i.e. ∑_*i*_ *μ_i_* = 1. It is assumed that all admixed samples included in the same analysis have similar ancestry proportions. If there are subgroups of admixed individuals with differing demographic histories, each subgroup can be analyzed separately.

Given a target haplotype, the prior probability for the state at any position is defined as follows. First the ancestry *i* is selected according to the probabilities *μ_i_*. Then the reference panel *j* is chosen according to the probabilities *p_ij_*. Finally, the donor haplotype is chosen randomly from the reference haplotypes for panel *j*. If *h* is from panel *j*, we write *q_ih_* = *P_ij_*/*n_j_* for the probability that the reference haplotype is *h* when the ancestry is *i*. Thus the prior probability that the state is (*i, h*) is *π*(*i, h*) = *μ_i_q_ih_*.

The parameter *T* is the number of generations since admixture. The distances between consecutive pairs of crossovers arising in the last *T* generations are exponentially distributed with mean 1/*T* Morgans (100/*T* centiMorgans).

Any crossover in the past *T* generations may change the ancestry state. Consider two markers indexed by *m* – 1 and *m* and separated by an interval of *d_m_* Morgans. The probability of at least one such crossover occurring in this interval is 1 – *e*^−*d_m_T*^. When a crossover occurs, a new ancestry is chosen according to the global ancestry probability vector ***μ***. The probability of a transition from ancestry state *i* to ancestry state *i′* is thus (1 – *e^−d_m_T^*) for *i* ≠ *i′*.

Changes in the donor reference haplotype *h* can occur regardless of whether there is a change in the ancestry. If there is no change in ancestry between the two markers, selection of a new donor reference haplotype occurs with probability 1 – *e^−d_m_ρ_i_^*, where *ρ_i_* is a population-specific switch rate and *i* is the ancestry state at both markers. If there is a change to ancestry *i′* between the two positions, a new donor reference haplotype *h* is always selected. In either case, the donor reference haplotype in the new state (*i′, h′*) is selected according to the probabilities *q_i′h′_*.

The resulting probability *P*(*S_m_* = (*i′, h′*)|*S*_*m*–1_ = (*i, h*)) of transitioning from state (*i, h*) to state (*i′, h′*) is given in Supplementary Methods 1.

If the target haplotype’s ancestry is *i* and the donor haplotype is from reference panel *j*, the emitted allele is the donor haplotype’s allele with probability 1 – *θ_ij_*, and is a different allele otherwise. The *θ_ij_* are miscopy rates which model recent mutation, genotype error, and gene conversion.

Let *I_m_*(*h*) = 1 if the allele at marker *m* on haplotype *h* matches the observed allele on the admixed haplotype, and let *l_m_*(*h*) = 0 otherwise. The emission probability *e_m_*(*i, h*) (probability of the data given the HMM state) for the allele at marker *m* on the admixed haplotype when the state is (*i, h*) and the reference haplotype *h* is from reference panel *j* is

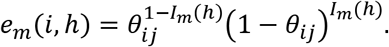

Given the parameter values and genetic data, we calculate the posterior probability of ancestry at each marker using hidden Markov model methods, which are described in Supplementary Methods 2. The assigned ancestry is the ancestry with the highest posterior probability.

### Estimating parameter values

A user can optionally specify parameter values. If not specified, values for the reference panel probabilities *p_ij_* and the within-ancestry switch rates *ρ_i_* are estimated from the reference panels and other parameters are assigned default values as described in Supplementary Methods 3. If the Expectation-Maximization (EM) option is turned on (the default), the values of ***μ*** and *T* will be updated based on several iterations of the estimation scheme described in Supplementary Methods 3. In the analyses presented in this paper, we use the default initial parameter values and perform EM updating.

If the user wishes to parallelize their analyses by chromosome, we recommend that the user run one autosomal chromosome first, and then use the output model file to specify the analysis parameters for other autosomes. This will reduce computing time and ensures consistency across chromosomes.

### Computational techniques

Many computational techniques have been developed that substantially reduce the computation time and memory requirements for genotype imputation. We incorporate several of these techniques in our local ancestry inference method. These techniques include a compressed representation for reference haplotypes in memory,^24; 29^ the use of a small custom panel of composite reference haplotypes for each admixed haplotype,^23; 25; 26^ and checkpointing of HMM backward probabilities.^30; 31^

The accuracy of local ancestry inference increases significantly with reference panel size (Figure 1), however large reference panels also increase the computational burden. In genotype imputation the use of a small, custom subset of reference haplotypes for each individual can reduce computation time by one or more orders of magnitude with no loss in accuracy.^26^ We have developed a fast method for generating a custom chromosome-length reference panel composed of composite reference haplotypes.^26^ Each composite reference haplotype is a mosaic of reference haplotype segments that incorporates long identity-by-descent segments between the reference haplotypes and a target haplotype. We create a custom set of composite reference haplotypes for each target haplotype, and we record the source reference panel for each reference haplotype segment so that the appropriate transition and emission probabilities can be used.

**Figure 1:**
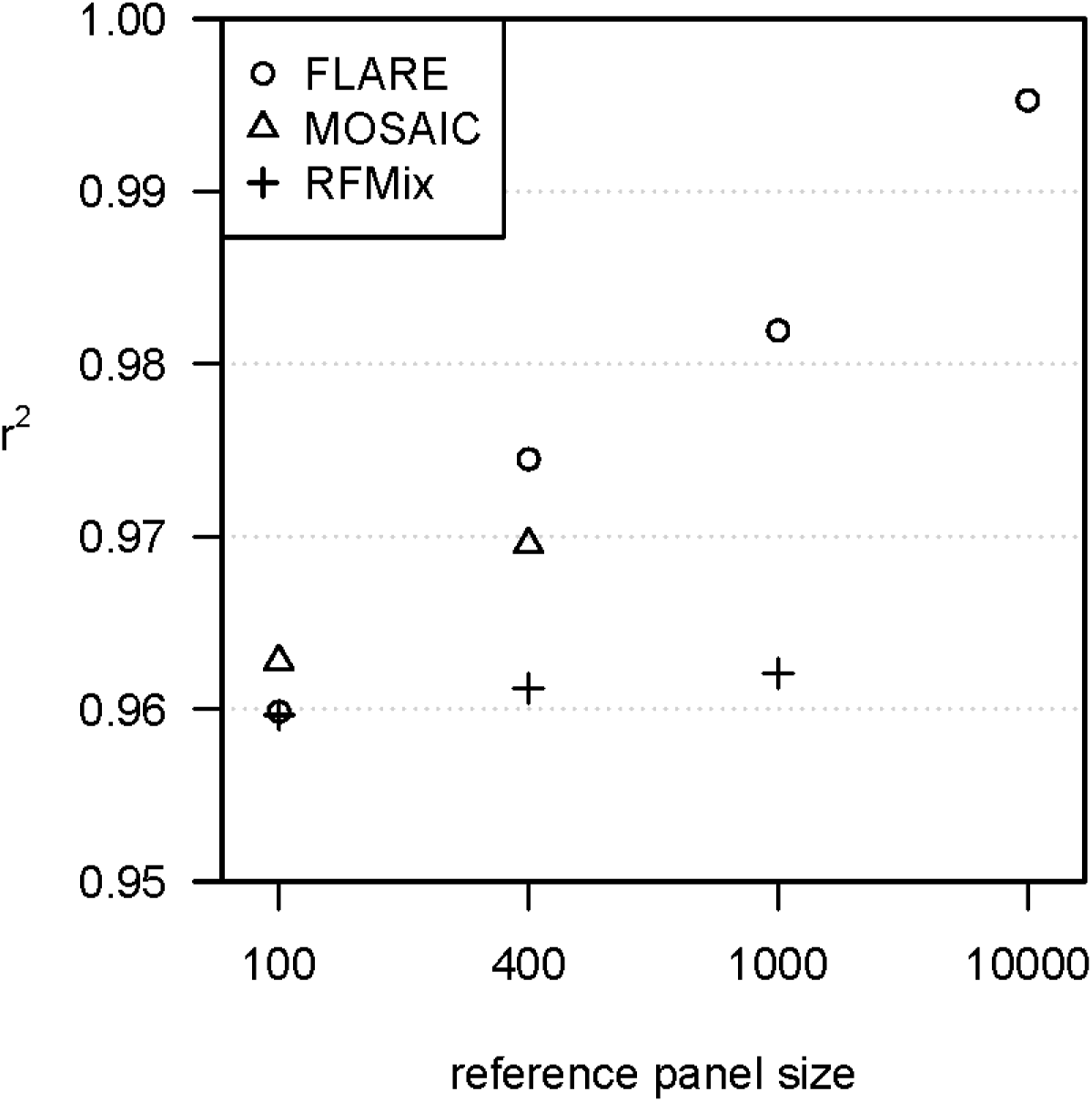
Accuracy when increasing reference panel size for simulated sequence data with three-way admixture. The y-axis shows squared correlation between true and inferred local ancestry dose averaged over ancestries (details in Methods). Each of the three ancestries is represented by a reference panel of size shown on the x-axis. Each analysis includes 400 admixed individuals, and results are averaged over four replicate simulations. MOSAIC could not analyze the data with 1000 or more individuals per reference panel within the available 256 GB of computer memory. RFMix could not analyze the data with 10,000 individuals per reference panel within the allotted five days.

We accommodate extremely large reference panels by compressing and storing the phased input data in bref3 format during program execution.^26^ This format compresses data for rare variants by storing the indices of haplotypes that carry each rare variant, and it compresses data for other variants by storing the distinct allele sequences in a genomic interval together with an array that maps each haplotype index to its allele sequence.^26^ The bref3 format enables an entire chromosome of reference and target haplotypes to be stored in memory and permits rapid lookup of haplotype alleles.

Checkpointing reduces the memory for HMM calculations for *M* markers from *O*(*M*) to 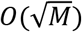 by storing forward probabilities at 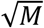 checkpoints, and re-calculating backward probabilities from the nearest preceding checkpoint when required.^30; 31^ Since there can be more than a million markers on a chromosome, checkpointing can produce a 1000-fold reduction in the memory required for HMM calculations, at the cost of a two-fold increase in computation time.

### Simulated data

We simulated genetic data from human out-of-Africa demographic models for three-way and four-way admixture, using modified versions of the human AmericanAdmixture_4B11 and OutOfAfrica_4J17 demographic models implemented in stdpopsim v0.1.2.^32^

The three-way model^11^ extends a model of African, European, and Asian demographic history^33^ to simulate admixture occurring 12 generations ago. The admixed population has 1/6 African, 1/3 European, and 1/2 Asian ancestry, an initial size of 30,000, and a growth rate of 5% per generation. We added population growth in the 10 most recent generations to the unadmixed populations, at rates of 19.3% (African population), 10.8% (European population) and 7.8% (Asian population), so that each population grows to approximately 100,000 individuals in order to permit sampling of large reference panels from these populations. We sampled 50,000 individuals from each of the three reference ancestries and 10,000 admixed individuals.

The four-way model extends the demographic history of African, European, Han Chinese, and Japanese populations inferred by Jouganous et al.^34^ We added an admixture event occurring 12 generations ago. The admixed population has 15% African, 15% European, 30% Chinese, and 40% Japanese ancestry, an initial size of 30,000, and a growth rate of 5% per generation. We sampled 400 individuals from each of the four reference ancestries and 400 admixed individuals.

We used SLiM^35^ (v3.7.1) for forward simulation of at least the most recent 10 × *N_e_* generations, followed by simulation of earlier generations with msprime^36^ (v1.1.1) to ensure full coalescence^37^. We simulated one full chromosome with characteristics similar to human chromosome 22. The simulated chromosome had a length of 51.3 Mb, a constant recombination rate of 1.44 × 10^−8^ per base pair per generation to match the average recombination rate of chromosome 22 as implemented in stdpopsim,^32^ and mutations at a rate of 1.44 × 10^−8^ per base pair per generation.^34^ During forward simulation, gene conversion was added at a rate of twice the base recombination rate and with a mean tract length of 300bp.

We constructed multiple data sets with a varying number of sampled individuals and different marker ascertainment schemes. The genetic data for each analysis were generated in three steps: 1) simulation of full demographic history and admixture 2) site ascertainment 3) analysis-specific data filtering. In the first step, the genotype data were simulated as described above. In the second step, two distinct ascertainment schemes were applied to produce simulated sequence data and simulated array data. For the array data, we removed all sites with a mean minor allele frequency (MAF) of less than 0.05 in the combined reference populations. For the sequence data, we removed all singletons. In the third step, the data sets were further filtered. After selecting individuals for a specific analysis, variants that were now singletons were removed. If the array data had more than 20,000 sites, 20,000 randomly selected sites were retained. If the sequence data had more than 1,000,000 sites (600,000 with reference panels of size 50,000), then 1,000,000 randomly selected sites were retained (600,000 with reference panels of size 50,000). Next, we added genotype error at a rate of *ε* = 0.0002, except at sites with MAF< 2ε where the error rate was set to MAF/2. Finally, all individuals (reference and admixed) were phased together with BEAGLE 5.2.

For each demographic history, we conducted four independent simulations and applied array and sequence ascertainment to each. This allowed four fully independent replicate analyses for each scenario, with no overlapping individuals or sites.

We inferred local ancestry with FLARE (v0.1.0), RFMix (v2.03-r0), and MOSAIC (v1.3.9). All programs were supplied with the same phased genotype data, genetic map, and reference panels. Parameters affecting the statistical analyses of the programs were kept at default values, except as noted. FLARE was run with posterior probabilities turned on (probs=true) since these were used to assess accuracy and calibration. For RFMix, 5 EM iterations were requested (-e). For MOSAIC, the number of grid points per centiMorgan (-GpcM) was set to the product of 0.0012 and the number of sites in the analysis. We found that this setting allowed MOSAIC to accurately analyze simulated sequence data. If a program didn’t report local ancestry at a site, the local ancestry of the closest preceding site was used.

The accuracy of each method was assessed with Pearson’s *r*^2^ by comparing the inferred and true local ancestry. True local ancestry was defined to be the population of residence of the ancestral chromosome segment 20 generations prior to sampling (8 generations prior to admixture). For each ancestry, we summed the local ancestry posterior probabilities for the two haplotypes to obtain an estimated diploid ancestry dose at each site, and we counted the number of copies of the ancestry in the true local ancestry (0, 1 or 2) to obtain the true diploid ancestry dose. We calculated the squared Pearson correlation of the estimated and true diploid ancestry dose across all individuals and sites. Separate *r*^2^ values were calculated for each ancestry and overall reported values are the unweighted mean *r*^2^ across all ancestries.

Each method reports posterior probabilities, which may be more or less well calibrated. For example, ideally, 90% of sites assigned 90% posterior probability of having ancestry 1 should actually be ancestry 1 and 10% should be another ancestry. Since the simulated data are statistically phased before inferring local ancestry, we cannot check calibration at the haplotype level, but must instead work at the diplotype level. Ideally, the average true ancestry dose for sites with an estimated diploid ancestry dose of 1.8 should be 1.8. To assess the calibration, we divide the range of possible estimated diploid ancestry doses into bins, and obtain the average true dose for each bin.

All analyses were run on a 20 core 2.4 GHz server with 256 GB memory and were run with 20 computational threads. We provide a repository containing all code for the generation and analysis of the simulated data presented here (see Web Resources).

### 1000 Genomes and Human Genome Diversity Panel data

We downloaded high-coverage sequence data for chromosome 1 from the Human Genome Diversity Project (HGDP) and from the 1000 Genomes Project (see Web Resources).^16^; ^17^ We merged the two data sets and excluded variants that were not biallelic SNPs with < 1% missingness and at least 5 copies of the minor allele in the combined data. After filtering, 2,021,066 SNPs remain on chromosome 1. We phased the data using Beagle 5.2 with the HapMap GRCh38 map (see Web Resources).^25^

We used the HGDP data for reference panels, assigning panels using the regional labels provided by the HGDP but omitting Oceania due to its smaller size and lack of relevance for the 1000 Genomes data. The panels range in size from 61 (America) to 223 (East Asia). We used FLARE with default settings to infer local ancestry in the 26 populations of the 1000 Genomes project, using a separate analysis for each of these populations. Ancestry proportions were obtained by averaging ancestry calls across sites and individuals.

## Results

### Simulated data

Figure 1 shows accuracy results for the three-ancestry simulation with sequence data. With small reference panel sizes (100 per ancestry) the three methods have similar performance (*r*^2^ between 0.960 and 0.963 for all three methods). With larger sample sizes, FLARE is most accurate (*r*^2^ = 0.995 with 10,000 individuals per reference panel) and RFMix is least accurate. For all methods we see an increase in accuracy with increasing reference panel size. This increase is greatest with FLARE and least with RFMix.

With simulated array data instead of sequence data, *r*^2^ accuracy decreases for FLARE and MOSAIC as expected, however *r*^2^ accuracy increases for RFMix (Figure S1). This suggests that the default settings of RFMix are better suited to array data than to sequence data, and that one should either thin the markers or use adjusted setting when analyzing sequence data with RFMix. Despite RFMix’s improved performance in array data, FLARE still has higher *r*^2^ accuracy than RFMix for larger reference panel sizes (0.977 vs 0.971 when there are 1000 individuals per reference panel).

Simulation studies typically employ reference panels of equal size for each ancestry, whereas in real analyses, some ancestries typically have fewer reference individuals. We thus investigated the accuracy when reference panels have unequal sizes. We found that all three methods performed well, with RFMix having the most consistently good performance (Figure S2).

We used the four-ancestry model to assess the ability of the methods to infer within-continent ancestry. For the sequence data, we find that FLARE has good resolution to distinguish all four ancestries (*r*^2^ = 0.954), whereas RFMix’s accuracy is severely reduced (*r*^2^ = 0.857), with most of this reduction being driven by the two Asian populations (Figure 2). MOSAIC was excluded from this comparison because it could not analyze the simulated four-ancestry sequence data within the available 256 GB of computer memory. For the array data, FLARE still performs the best (*r*^2^ = 0.941), but RFMix (*r*^2^ = 0.934) and MOSAIC (*r*^2^ = 0.921) are only slightly less accurate in this case (Figure S3).

**Figure 2:**
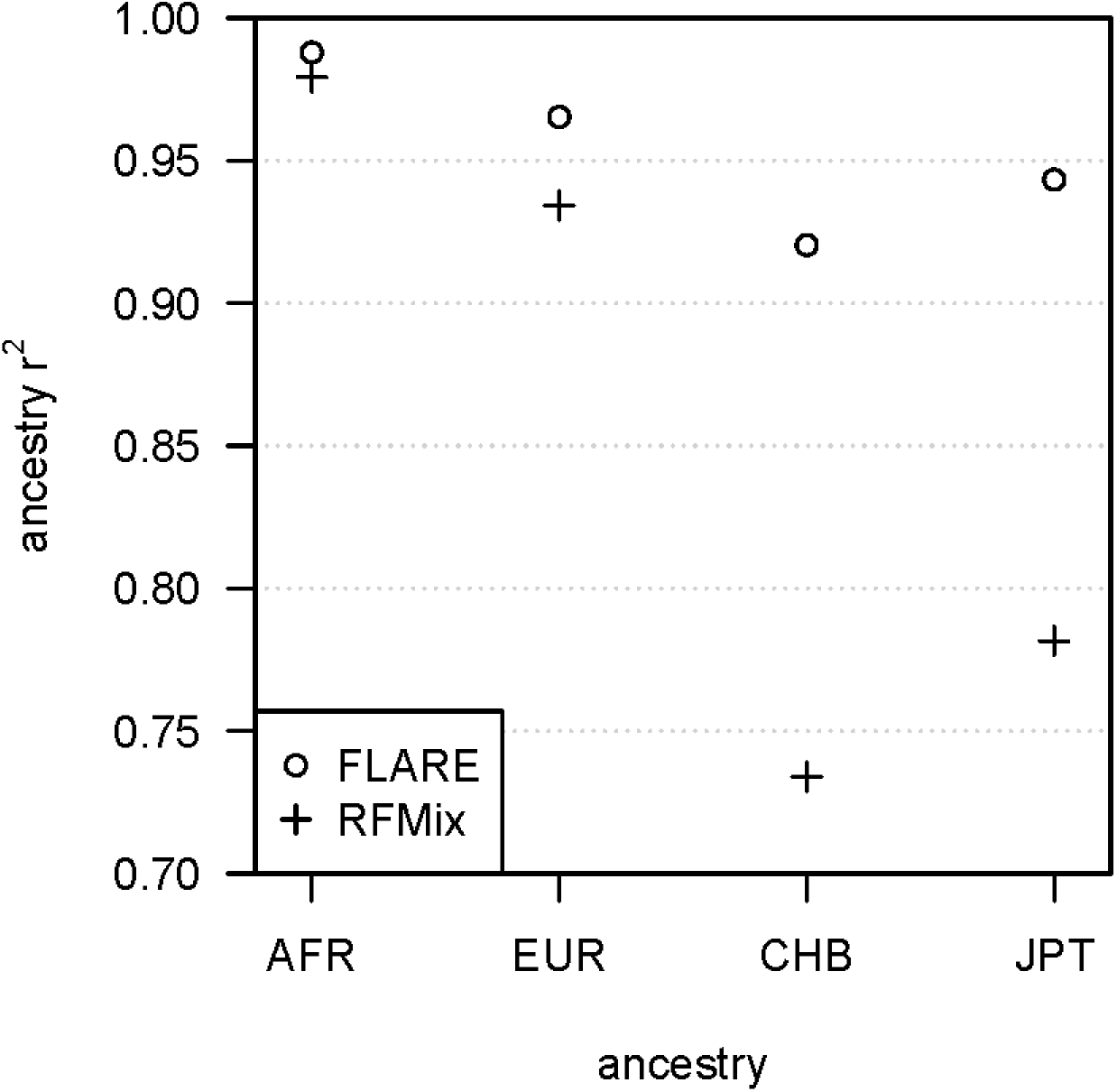
Accuracy by ancestry for simulated sequence data with four-way admixture. The y-axis is the squared correlation between the true and inferred ancestry dose for a single ancestry. The ancestry is shown on the x-axis (AFR is African, EUR is European, CHB is Han Chinese from Beijing, JPT is Japanese from Tokyo). The simulated sequence data have 400 admixed individuals and 400 individuals in each of the four reference panels. Results are averaged over four replicate simulations. MOSAIC could not analyze these data within the available 256 GB of computer memory.

We found that both FLARE and MOSAIC are well calibrated in terms of their reported posterior ancestry probabilities (Figure S4). In contrast, RFMix’s output probabilities are not well calibrated. This is not too surprising given the generative probabilistic modelling that underlies FLARE and MOSAIC, in contrast to the discriminative approach taken by RFMix.

FLARE has much faster computation times than the other two methods (Figure 3). Compute times for FLARE are relatively insensitive to the reference panel size due to its use of composite reference haplotypes (see Methods). FLARE’s analyses of 400 admixed individuals in the three-way admixture setting with sequence data take less than 5 minutes for reference panel sizes of up to 1000 per ancestry and an average of 10 minutes for 10,000 individuals in each of the three reference panels. FLARE’s analysis of 10,000 admixed individuals with 50,000 individuals in each of the three reference panels took an average of 30 minutes.

**Figure 3:**
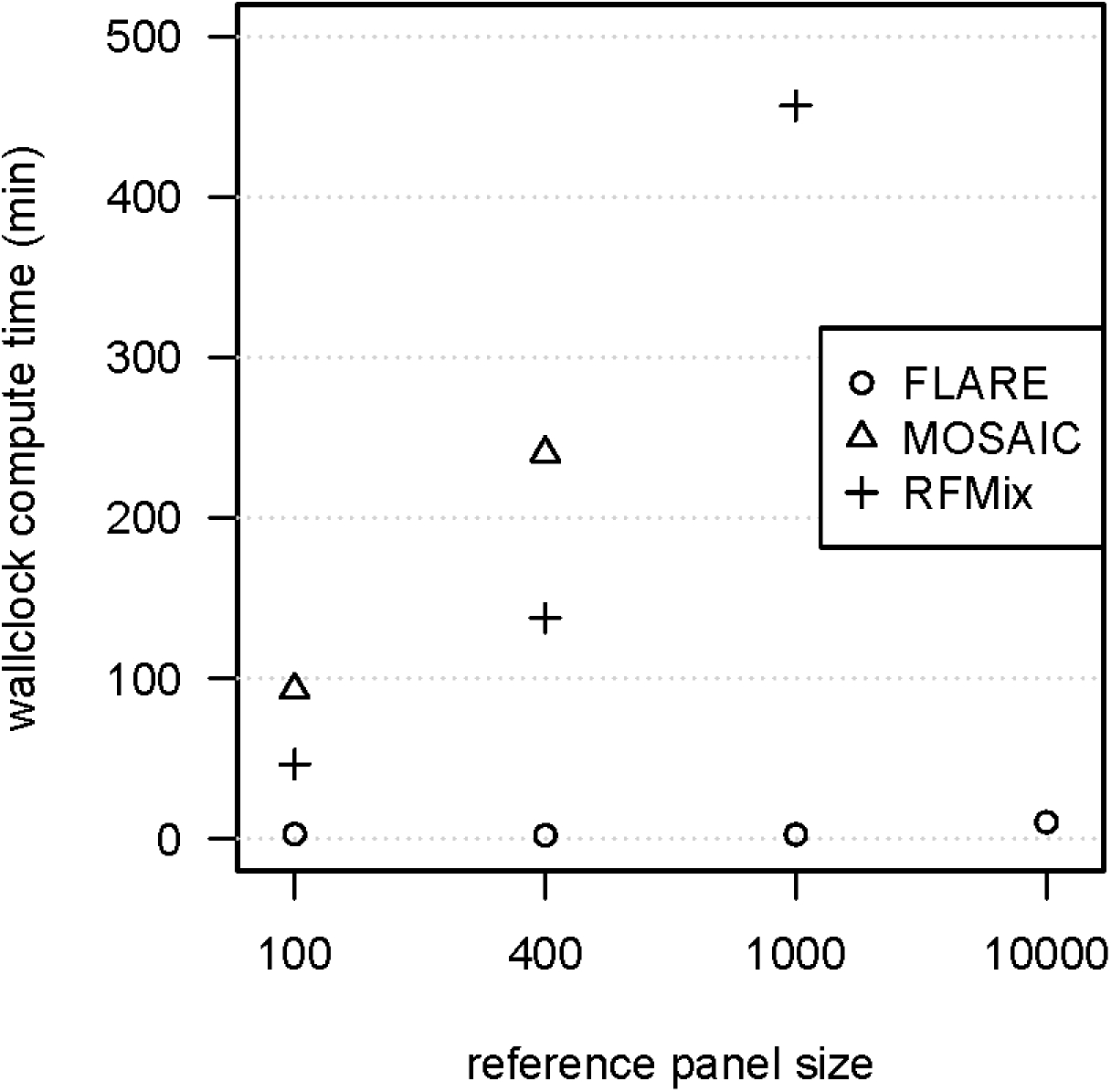
Computation time for simulated sequence data with three-way admixture. Wallclock computation time in minutes is shown on the y-axis. Reference panel size for each of the three ancestries is shown on the x-axis. Each analysis includes 400 admixed individuals, and results are averaged over four replicate simulations. Analyses of a simulated chromosome modeled on chromosome 22 were run on compute nodes with 20 cores. MOSAIC could not analyze the data with 1000 or more individuals per reference panel within the available 256 GB of computer memory. RFMix could not analyze the data with 10,000 individuals per reference panel within the allotted five days (7200 minutes).

For the four-way admixture with sequence data, 400 admixed individuals, and 400 individuals in each of the four reference panels, FLARE took an average of 7 minutes, while RFMix took an average of 357 minutes. Compute times are expected to scale approximately linearly in the number of admixed individuals for each of the three methods.

### 1000 Genomes local ancestry analysis

We inferred local ancestry for each of the 26 populations of the 1000 Genomes Project using 6 regional reference panels from the HGDP. Estimated ancestry proportions from this analysis are shown in Table 1. Results generally match expectation. For example, the unadmixed African populations are inferred to have 98-100% African ancestry. Native American ancestry originally derives from Siberia,^38^ which may partially explain the inferred East Asian ancestry in the admixed American populations, although post-colonial migration from Asia may also play a role.^39^ Finns (FIN) are inferred to have 4% East Asian ancestry, which is concordant with previous studies that have found evidence of an Asian contribution to the gene pool in Finns.^40; 41^

**Table 1.** Inferred ancestry proportions in 1000 Genomes Project chromosome 1 data for six ancestries using HGDP reference panels. Ancestry proportions > 20% are bolded’ proportions < 3% are grayed. Descriptions of the populations can be found in Supplementary Information Table 1 of the 1000 Genomes Project’s phase 3 paper.^43^ Recently admixed populations from the Americas are marked with an asterisk.

| Region | Population | African | East |  | Central/South |  | Middle Eastern |
| --- | --- | --- | --- | --- | --- | --- | --- |
|  |  |  | Asian | European | Asian | American |  |
| Africa | ACB* | **0.89** | 0.00 | 0.10 | 0.01 | 0.00 | 0.00 |
|  | ASW* | **0.77** | 0.01 | **0.19** | 0.00 | **0.03** | 0.00 |
|  | ESN | **1.00** | 0.00 | 0.00 | 0.00 | 0.00 | 0.00 |
|  | GWD | **0.99** | 0.00 | 0.01 | 0.00 | 0.00 | 0.00 |
|  | LWK | **0.98** | 0.00 | 0.00 | 0.00 | 0.00 | 0.02 |
|  | MSL | **1.00** | 0.00 | 0.00 | 0.00 | 0.00 | 0.00 |
|  | YRI | **1.00** | 0.00 | 0.00 | 0.00 | 0.00 | 0.00 |
| America | CLM* | 0.09 | 0.02 | **0.61** | 0.02 | **0.26** | 0.02 |
|  | MXL* | 0.06 | 0.03 | **0.43** | 0.02 | **0.45** | 0.01 |
|  | PEL* | 0.04 | 0.06 | **0.20** | 0.01 | **0.69** | 0.00 |
|  | PUR* | **0.16** | 0.01 | **0.65** | 0.02 | **0.14** | 0.02 |
| East Asia | CDX | 0.00 | **1.00** | 0.00 | 0.00 | 0.00 | 0.00 |
|  | CHB | 0.00 | **1.00** | 0.00 | 0.00 | 0.00 | 0.00 |
|  | CHS | 0.00 | **1.00** | 0.00 | 0.00 | 0.00 | 0.00 |
|  | JPT | 0.00 | **1.00** | 0.00 | 0.00 | 0.00 | 0.00 |
|  | KHV | 0.00 | **1.00** | 0.00 | 0.00 | 0.00 | 0.00 |
| Europe | CEU | 0.00 | 0.00 | **1.00** | 0.00 | 0.00 | 0.00 |
|  | FIN | 0.00 | 0.04 | **0.96** | 0.00 | 0.00 | 0.00 |
|  | GBR | 0.00 | 0.00 | **1.00** | 0.00 | 0.00 | 0.00 |
|  | IBS | 0.02 | 0.00 | **0.98** | 0.00 | 0.00 | 0.00 |
|  | TSI | 0.00 | 0.00 | **1.00** | 0.00 | 0.00 | 0.00 |
| South Asia | BEB | 0.00 | 0.00 | 0.00 | **1.00** | 0.00 | 0.00 |
|  | GIH | 0.00 | 0.00 | 0.00 | **1.00** | 0.00 | 0.00 |
|  | ITU | 0.00 | 0.00 | 0.00 | **1.00** | 0.00 | 0.00 |
|  | PJL | 0.00 | 0.00 | 0.00 | **1.00** | 0.00 | 0.00 |
|  | STU | 0.00 | 0.00 | 0.00 | **1.00** | 0.00 | 0.00 |

Our initial analyses of chromosome 1 with parameter estimation took 16.6 hours (38 mins per population on average). Analyses of other chromosomes that use the parameters estimated from the chromosome 1 data would use much less time. For example, when we repeated the chromosome 1 analysis using the parameters estimated in the first analysis, the second analysis took only 2.4 hours (6 minutes per population on average). The first and second analysis produced identical estimated ancestry probabilities. Parameter estimation is only needed for one autosomal chromosome, with analyses of the other autosomal chromosomes using the same parameters.

## Discussion

We have presented FLARE, a new method for local ancestry inference. We showed that FLARE has superior accuracy, computing efficiency, and scalability compared to RFMix and MOSAIC. FLARE was able to analyze data with 10,000 admixed individuals and three 50,000 member reference panels in ten minutes on a computer with 20 CPU cores, while RFMix was unable to complete analysis with 400 admixed individuals and three 10,000 member reference panels within five days on the same computer. MOSAIC was even more limited in the data that it could analyze due to memory constraints, and was significantly slower than RFMix. FLARE had the highest accuracy of the three methods except when reference panel sizes were small.

A notable feature of the results of the simulations studies is that FLARE can distinguish within-continent ancestry, such as distinguishing between Japanese and Chinese ancestry. In contrast, RFMix had difficulty distinguishing these ancestries. This suggests the potential to infer local ancestry in admixtures that are subtler than the continental-level admixtures that have previously been the focus of attention.

FLARE’s estimated posterior probabilities of ancestry are well calibrated. This is important when one wants to incorporate ancestral uncertainty in downstream analyses.

FLARE is a user-friendly java program with a command-line interface similar to Beagle.^25^ When there is a one-to-one matching of ancestries and reference panels, the only input data required by FLARE are phased reference and target VCF files,^42^ a genetic map file, and a file specifying the reference panel assignment for each reference individual. FLARE outputs a VCF file containing the input genotypes and phased local ancestry calls. As an option, the posterior local ancestry probabilities can also be included in the output VCF file. FLARE also outputs a model file giving the estimated parameters. The model can be used in future analyses of the same population to reduce computing time and ensure consistency between analyses.

FLARE’s user-friendly and robust software implementation, its computational speed and ability to scale to extremely large data sets, and its high accuracy make it a useful tool for local ancestry inference in the increasingly large and diverse genetic data that are being generated.

## Web Resources

1000 Genomes Project high-coverage sequence data: http://ftp.1000genomes.ebi.ac.uk/vol1/ftp/data_collections/1000G_2504_high_coverage/working/20190425_NYGC_GATK/

Human Genome Diversity Project high-coverage sequence data: ftp://ngs.sanger.ac.uk/production/hgdp/hgdp_wgs.20190516/

Beagle: http://faculty.washington.edu/browning/beagle/beagle.html

HapMap GRCh38 map: http://bochet.gcc.biostat.washington.edu/beagle/genetic_maps/plink.GRCh38.map.zip

stdpopsim: https://github.com/popsim-consortium/stdpopsim

msprime: https://github.com/tskit-dev/msprime

MOSAIC: https://maths.ucd.ie/~mst/MOSAIC/

RFMIX: https://github.com/slowkoni/rfmix

Simulation and analysis pipeline: https://github.com/rwaples/lai-sim

FLARE: https://github.com/browning-lab/flare

## Acknowledgements

Research reported in this publication was supported by the National Human Genome Research Institute of the National Institutes of Health under award numbers HG010869 and HG008359. The content is solely the responsibility of the authors and does not necessarily represent the official views of the National Institutes of Health.

---

**Figure S1:**
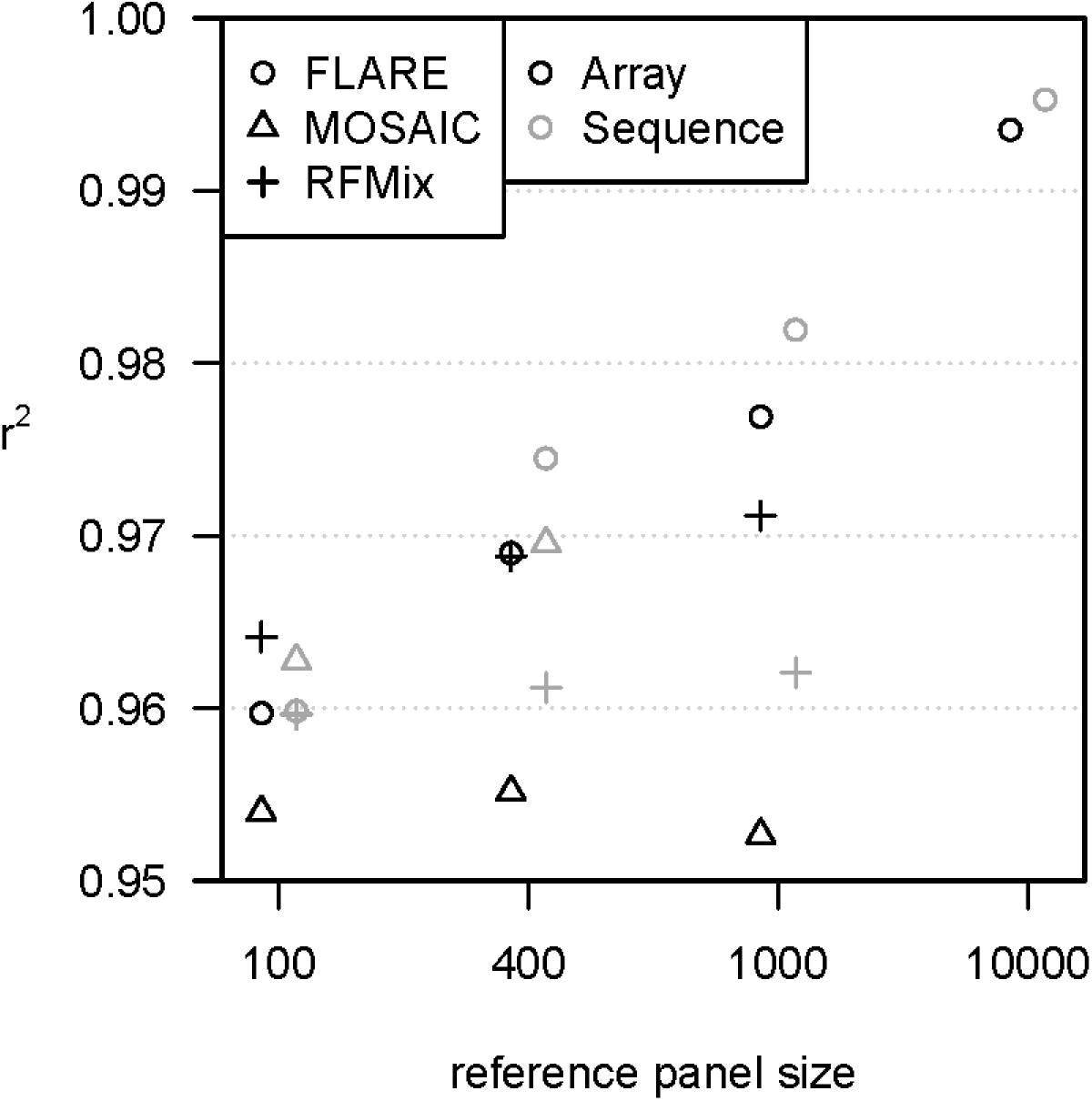
Accuracy when increasing reference panel size for simulated array data with three-way admixture. Results from Figure 1 for sequence data are shown in gray. The y-axis shows squared correlation between true and inferred local ancestry dose averaged over ancestries (details in Methods). Each of the three ancestries is represented by a reference panel of size shown on the x-axis. Each analysis includes 400 admixed individuals, and results are averaged over four replicate simulations. MOSAIC could not analyze the sequence data with 1000 or more individuals per reference panel or the array data with 10,000 individuals per reference panel within the available 256 GB of computer memory. RFMix could not analyze the data with 10,000 individuals per reference panel within the allotted five days.

**Figure S2:**
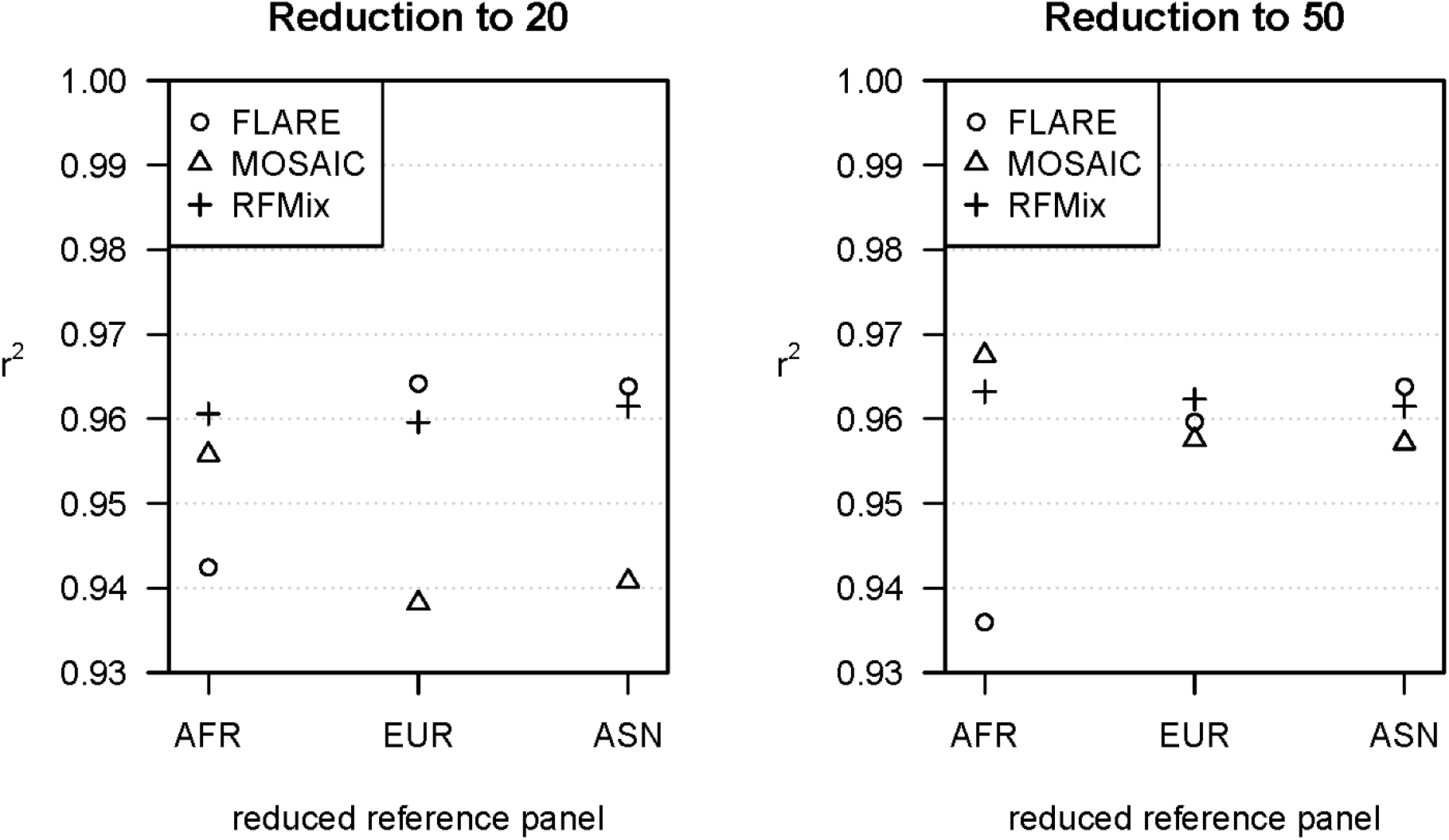
Accuracy when reducing the size of one of three reference panels. The data are simulated sequence data for three-way admixture. Each reference panel has size 400, except for the reference panel that is denoted on the x-axis (AFR is African, EUR is European, ASN is Asian) which has size 20 (left plot) or 50 (right plot). The y-axis shows squared correlation between true and inferred local ancestry dose averaged over ancestries (details in Methods). Each analysis includes 400 admixed individuals, and results are averaged over four replicate simulations.

**Figure S3:**
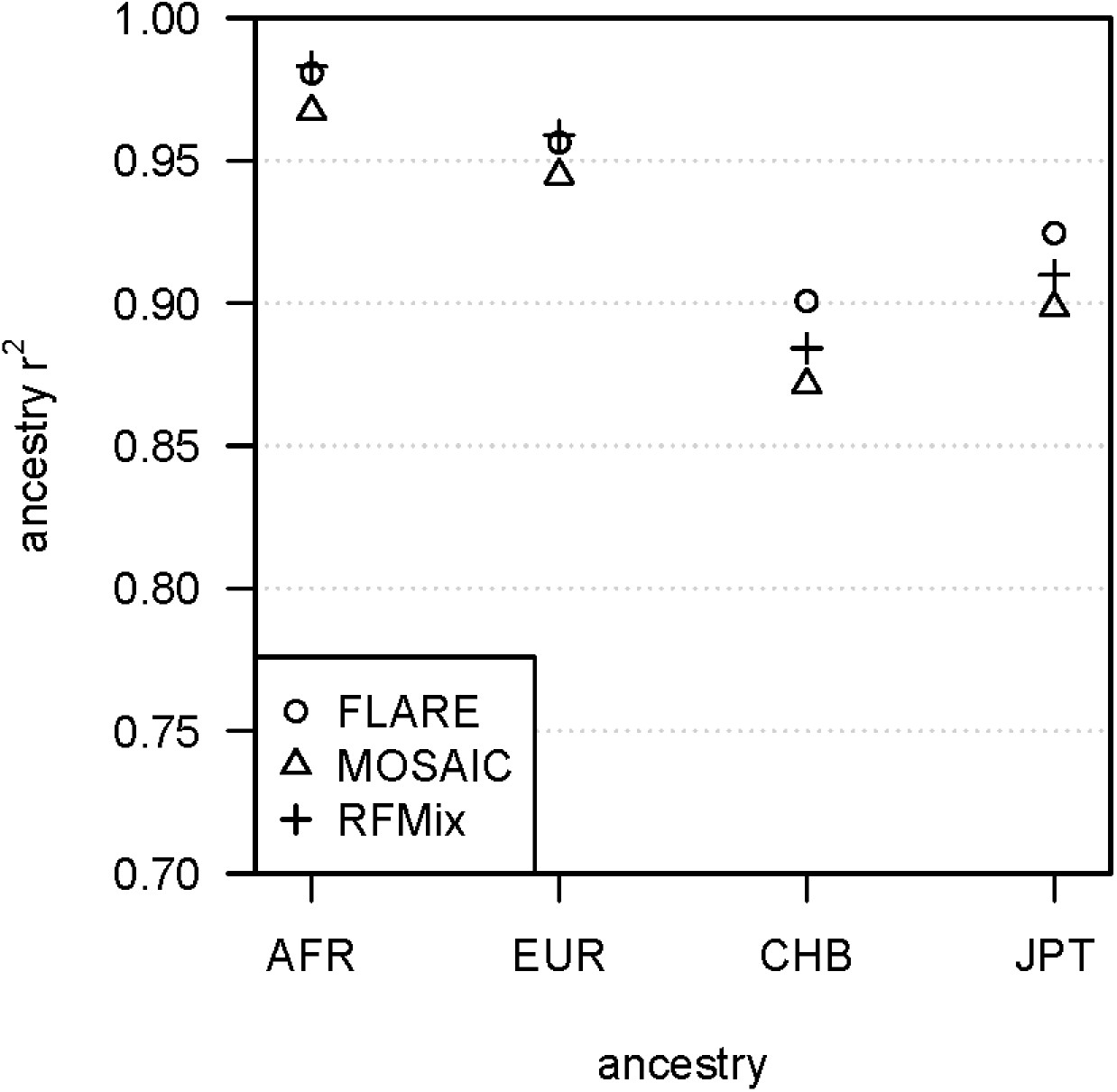
Accuracy by ancestry for the simulated array data with four-way admixture. The y-axis is the squared correlation between the true and inferred ancestry dose for a single ancestry. The ancestry is shown on the x-axis (AFR is African, EUR is European, CHB is Han Chinese from Beijing, JPT is Japanese from Tokyo). The simulated array data have 400 admixed individuals and 400 individuals in each of the four reference panels. Results are averaged over four replicate simulations.

**Figure S4:**
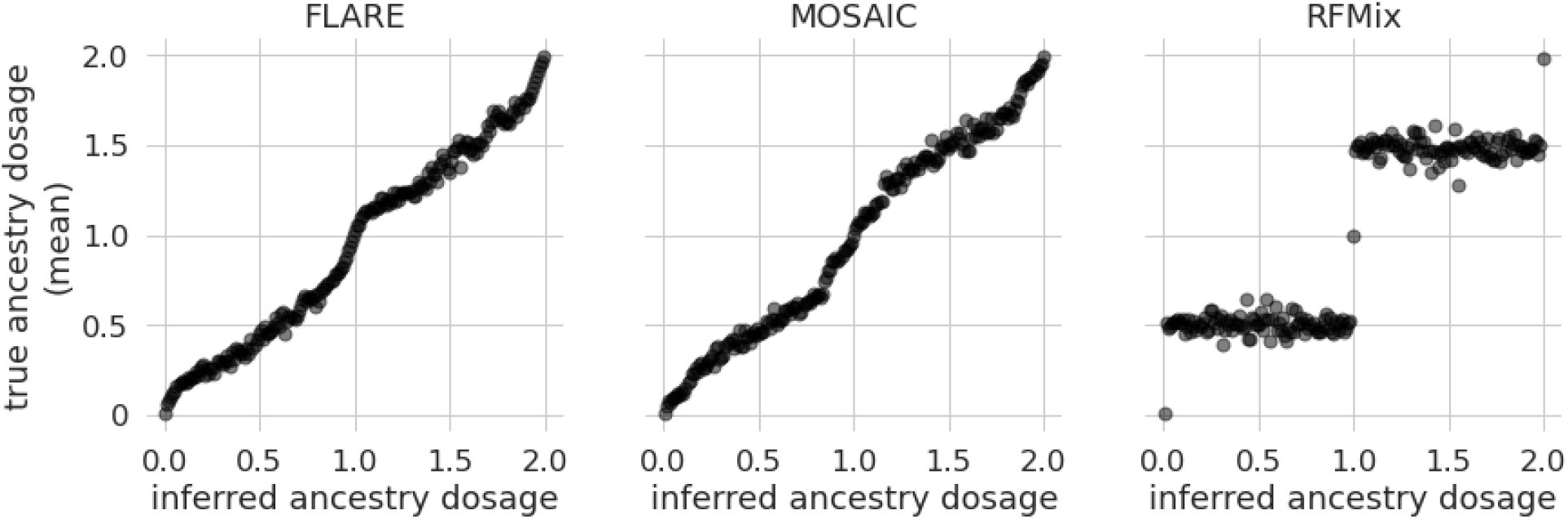
Calibration of estimated diploid ancestry dose on simulated data. Estimated diploid ancestry dose is binned into bins of width 0.01 along the x-axis. The y-axis is the average true diploid ancestry dose for each bin. Results for FLARE, MOSAIC, and RFMix are shown in the left, middle, and right panels respectively. All simulation scenarios for which all three methods successfully completed are included in these plots.

## Supplementary Methods 1: Transition probabilities

The transition probabilities described in the main text can be expressed as:

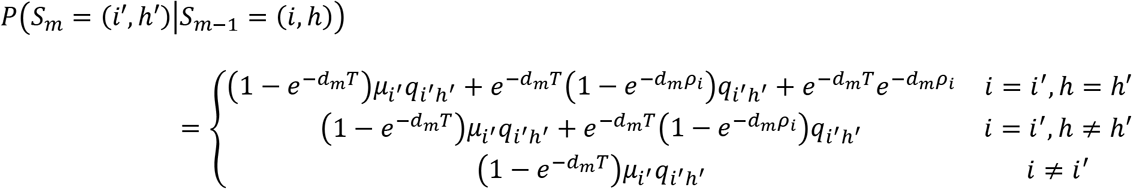

## Supplementary Methods 2: Algorithm for posterior probabilities of ancestry

We estimate the posterior ancestry probabilities using the hidden Markov model forward-backward algorithm.^44^

Consider an admixed haplotype, ***Y***. Let *Y_m_* be the allele at marker *m*, with markers indexed 1,…, *M*. The forward probabilities are

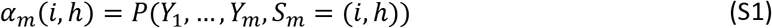

where *S_m_* represents the (ancestry, haplotype) state at the *m*th marker. The backward probabilities are

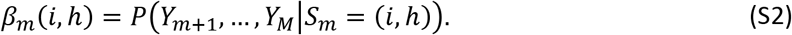

### Forward probabilities at first marker

For each ancestry *í* and haplotype *h*,

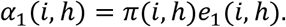

where *π*(*i, h*) is the prior probability that the state is (*i, h*), and the emission probability *e_m_*(*i, h*) is the probability of observing the allele *Y_m_* at marker *m* on the admixed haplotype when the hidden state at this marker is (*i, h*).

### Forward probabilities

Suppose we have already calculated *α*_*m*–1_(*i, h*) for all (*i, h*), and we want to calculate *α_m_*(*i′, h′*). Let *d_m_* be the distance in Morgans between markers *m* – 1 and *m*. Pre-calculate

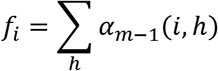

for each *i*, and

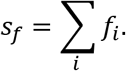

The values *f_i_* and *S_f_* are temporary variables that are over-written for each successive marker. Their purpose is to avoid duplicate calculation.

Then for each *i′* and *h′* calculate (using equation S1)

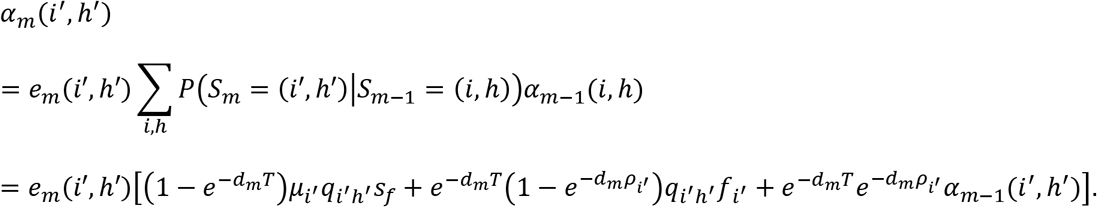

In the computation, we normalize the *α_m_*(*i′, h′*) to sum to one and store the normalization factors in order to avoid numerical underflow.

### Backwards probabilities

Let *β_M_*(*i, h*) = 1 for all ancestries *i* and reference haplotypes *h*.

Suppose the *β*_*m*+1_(*i, h*) values have been calculated for all ancestries *i* and reference haplotypes *h*. Let *d*_*m*+1_ be the distance in Morgans between markers *m* and *m* + 1. Pre-calculate

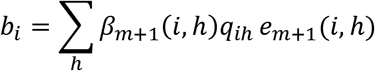

for each *i*, and *s_b_* = ∑*^i^b_i_μ_i_*

The values *b_i_* and *s_b_* are temporary variables that are over-written for each successive marker. Their purpose is to avoid duplicate calculation.

Then for each *i* and *h*, calculate (using equation S1)

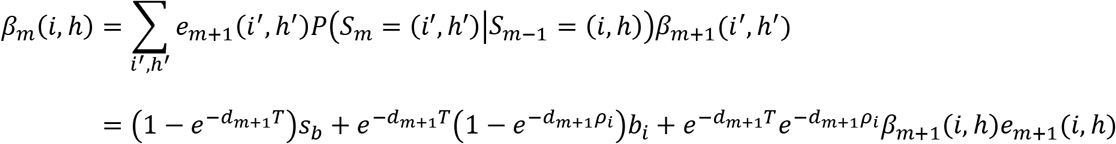

In the computation, we normalize the values of *β_m_*(*i, h*) to sum to one and store the normalization factors to avoid numerical underflow.

### Posterior probability of ancestry

Let

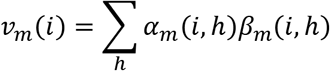

The posterior probability of ancestry *i* at marker *m* is *w_m_*(*i*) = *v_m_*(*i*)/∑*_i′_v_m_*(*i′*).

## Supplementary Methods 3: Initialization and updating parameter values

The initial values of the parameters are set as described below, or as specified by the user. If the EM updating option is turned on (which it is by default), each EM iteration estimates local ancestry for 100 randomly selected admixed haplotypes and the ancestry proportions and admixture time are updated as described below. Twenty EM iterations are performed unless the EM updating converges sooner. Convergence is defined as a relative change less than 5% in each ancestry proportion *μ_i_* from the value in the preceding iteration, excluding those ancestries for which *μ_i_* < 0.001

### Miscopy probabilities *θ_i,j_*

The default miscopy probabilities are the same for each ancestry and panel, and are defined as: *θ_i,j_* = *λ*/(2*λ* + 2*N*) where *λ* = 1/(log*N* + 0.5) and *N* is the total number of reference haplotypes.^45^ We do not update this parameter.

### Panel probabilities *p_ij_* and switch rates *ρ_i_*

The panel probabilities are obtained via training on the reference panel. Considering ancestry *i*^*^, which is represented by one reference panel, we take one haplotype at a time out of that reference panel and run the forwards-backwards algorithm using all other reference haplotypes. For this analysis we set *μ_i^*^_* = 1, *μ_i_* = 0 for *i* ≠ *i*^*^, *T* = 0, and *p*_*i*^*^*j*_ = *n_j_*/*N* where *n_j_* is the number of reference haplotypes in panel *j*. We use the default miscopy probabilities *θ_ij_* defined in the preceding section, and we set *ρ_i_* = 4*N_e_/N* where *N_e_* = 50,000.^24; 45^ We perform the analysis for 100 haplotypes selected at random from the reference panel.

The updated panel probability is the average posterior probability that the copied haplotype is from panel *j*, given that the ancestry is *i*. The posterior probability for state (*i, h*) is proportional to *α_m_*(*i, h*)*β_m_*(*i, h*). The selected haplotypes are indexed by *k*.

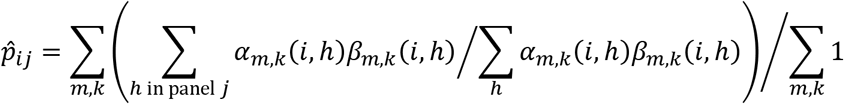

The updated switch rate *ρ_i_* is determined from the posterior probabilities of a change of haplotype state, as follows:

The probability of transitioning to the same state is:

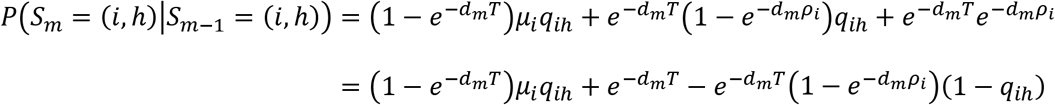

Solving for (1 – *e*^−*d_m_ρ_i_*^) gives:

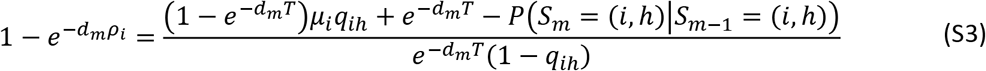

We write *τ_m,i_* = 1 – *e^−d_m_ρ_i_^*. We estimate *τ_m,i_* using the observed transition probabilities in place of the prior transition probabilities *P*(*S_m_* = (*i, h*)|*S*_*m*–1_ = (*i, h*)):

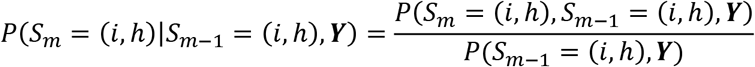

We average over haplotype state *h*, weighting by the observed state probabilities conditional on ancestry *i*,

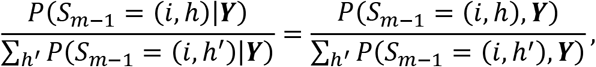

in the right-hand side of equation S3 to obtain:

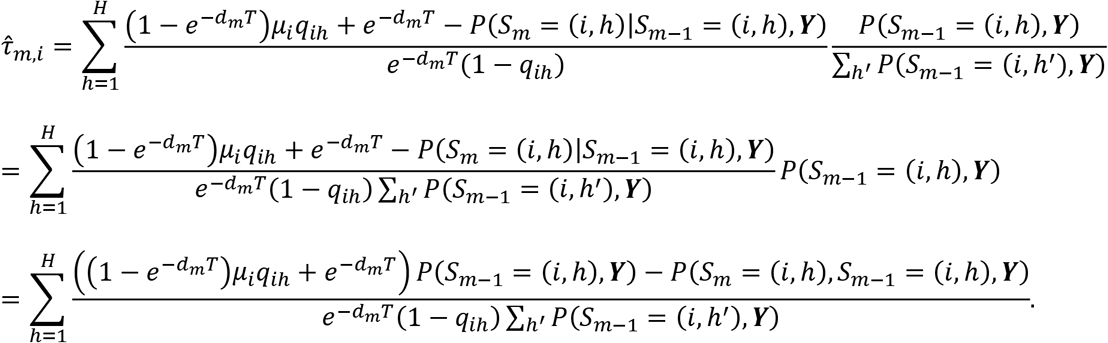

At each marker *m* > 1,

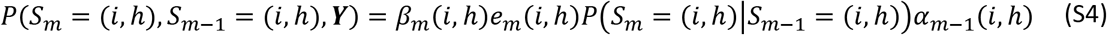

and

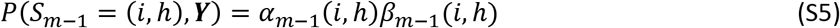

After we have estimated the 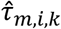 for each marker *m* and each target haplotype *k*, we estimate the constant of proportionality *τ* in the relationship 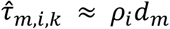 a slope estimator weighted by the conditional probability of ancestry *i* given the data, ∑*_h_P*(*S*_*m*–1_ = (*i,h*)|***Y***):

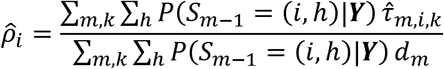

Note that

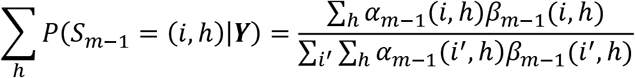

After initializing the *p_ij_* and *ρ_i_*, these parameters are fixed for the remainder of the analysis.

### Ancestry proportions, *μ_i_*

The default initial value is 1/*A*, where *A* is the number of ancestries. The updated value following each EM iteration is a weighted average of the posterior probability *w_m_*(*i*) for ancestry *i*. We include only positions for which the posterior probability of the ancestry is at least 0.9 in order to speed convergence. The selected haplotypes are indexed by *k*.

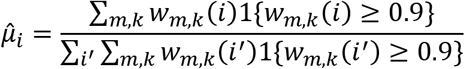

### Admixture time *T*

The default initial value of *T* is 10 generations.

The updated admixture time is determined from the posterior probabilities of a change of ancestry state, as follows:

The probability of transitioning to the same ancestry state is

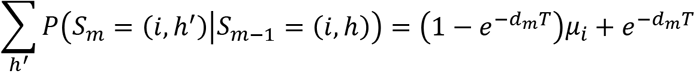

Solving for (1 – *e^−d_m_T^*) we obtain

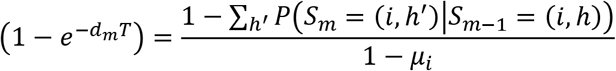

We write *γ_m_* = 1 – *e^−d_m_T^*. We estimate *γ_m_* using the observed transition probabilities in place of the prior transition probabilities *P*(*S_m_* = (*i, h*)|*S*_*m*–1_ = (*i, h*)):

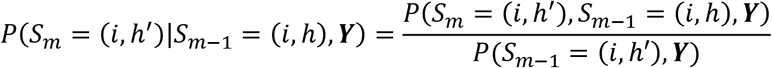

We average over haplotype state *h* and ancestry *i* at marker *m* – 1, weighting by the observed state probabilities:

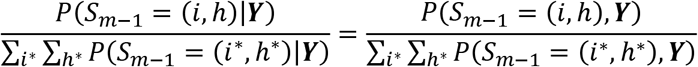

to obtain

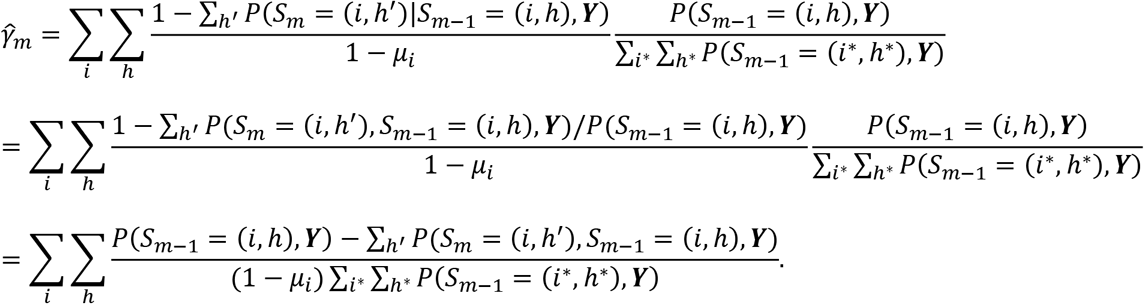

At each marker *m* > 1,

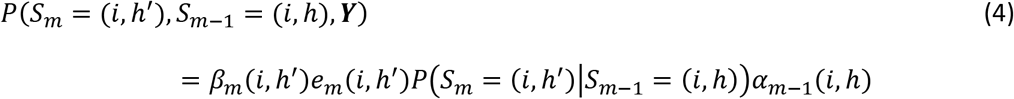

and

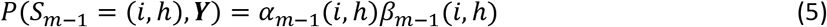

After we have estimated the 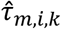 for each marker *m* and each target haplotype *k*, we estimate the constant of proportionality *T* in the relationship 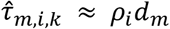 as:

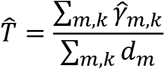

## Notes

### Competing Interest Statement

The authors have declared no competing interest.

